# Food-entrainment of circadian timekeeping in the dorsal vagal complex

**DOI:** 10.1101/2024.12.20.629643

**Authors:** Lukasz Chrobok, Charlotte Muir, Tanya Chonkria Kaur, Iliana Veneri, Timna Hitrec, Michael Ambler, Anthony Edward Pickering, Hugh David Piggins

## Abstract

The dorsal vagal complex (DVC) is a multi-component brainstem satiety centre which has gained attention as a key target of anti-obesity pharmacotherapies. Our recent studies revealed its circadian timekeeping properties, with molecular and electrophysiological 24h rhythms persisting independently of the primary hypothalamic clock. However, the factors entraining these brainstem oscillators, and the downstream transcriptional targets of the DVC molecular clock remain unclear. Here, using fluorescent *in situ* hybridisation, we demonstrate core clock gene expression in inhibitory and excitatory neuronal populations of the DVC, as well as in its output cholinergic vagal neurons. We further reveal that the molecular clock is associated with rhythmic expression of numerous neurotransmitter receptor genes in the DVC *in vivo*, with the phase of both clock and clock-controlled gene expression tightly regulated by meal timing. These findings uncover food-entrained circadian rhythms in the DVC and have important implications for clinical studies targeting brainstem satiety mechanisms.

## INTRODUCTION

Feeding is a vital behaviour for survival, and organisms have evolved complex mechanisms to ensure adequate food intake. In mammals, neural circuits that regulate feeding are finely tuned to nutrient composition and the daily patterns of food availability in the environment. This allows organisms to not only respond to food presence but also anticipate its availability ^1–3^.

The suprachiasmatic nucleus (SCN) of the hypothalamus is the central circadian clock in mammals, generating approximately 24-hour rhythms in physiology and behaviour. The SCN’s timekeeping ability is driven by the rhythmic expression of molecular clock components. These core clock genes operate within transcription-translation feedback loops (TTFL) to maintain a cycle close to 24 hours. Their protein products are transcription factors that in turn regulate the expression of numerous clock-controlled genes to ensure timely transcription and availability of proteins including signalling molecules, neurotransmitter receptors, and ion channels ^4,5^.

SCN neurons can adjust the phase of the molecular clock in response to environmental light, via their innervation by the retina, and thus constitute a light-entrainable oscillator ^6^. However, under time-restricted feeding (TRF), where food is provided at specific times, feeding becomes an additional, competing time cue (Zeitgeber). Evidence indicates that food anticipatory activity (FAA), the behavioural arousal preceding repetitive timed food presentation, operates largely independent of the SCN and is driven by a network of extra-SCN oscillators in the brain and peripheral organs, collectively termed the food-entrainable oscillator (FEO) ^7–10^. Despite extensive research, the FEO has not been localised to a single brain region or peripheral tissue, with several brain centres involved in feeding behaviour identified as contributors ^11^.

While many feeding centres are in the hypothalamus, we have identified that the dorsal vagal complex (DVC) in the hindbrain exhibits robust circadian timekeeping properties ^12^. The DVC is a crucial hub for ingestive, metabolic, cardiovascular, and other homeostatic functions, serving as both a primary source and recipient of vagal innervation ^13,14^. Known in the metabolic control field as the brainstem satiety centre, the DVC has recently gained attention for its fundamental role in the action of obesity medications ^15–17^. The DVC comprises three anatomically and functionally distinct structures, all expressing circadian rhythmicity: the area postrema (AP; the sensor of the complex), the nucleus of the solitary tract (NTS; the processor), and the dorsal motor nucleus of the vagus (DMV; the peripheral output). Additionally, we have recently found that non-neuronal ependymal cells lining the central canal and the fourth ventricle within the DVC rhythmically express molecular clock genes *ex vivo* ^12^. Despite clear evidence implicating the DVC in meal termination and more broadly in feeding behaviour, it is still unclear if and how food intake acts as a Zeitgeber for the DVC circadian oscillators.

Previous studies, including our own, have demonstrated that circadian timekeeping in the DVC of mice and rats is highly sensitive to diet composition ^18–20^. The aim of this study was to evaluate whether the DVC’s circadian properties are similarly sensitive to the timing of food intake and to examine the consequences of altered feeding schedules on gene expression within this brainstem complex. By investigating competing Zeitgeber conditions, our experiments show that the DVC’s circadian rhythms are responsive to the timing of food intake, rather than light-dark cycles. These food-entrained rhythms in components of the molecular clock further shape the rhythmic transcription of clock-controlled genes, thus aligning DVC gene expression to feeding behaviour.

## RESULTS

### Neurochemical Profile of Clock Gene-Expressing Cells in the Dorsal Vagal Complex (DVC)

In the SCN, the primary circadian clock, there is functional delineation between cell populations expressing clock genes. These functional groups have distinct neurochemical identities and spatiotemporal organisation within the SCN ^21,22^. However, the neurochemical identity of clock gene-expressing cells within the DVC remains unexplored and it is unclear whether there is spatiotemporal organisation to its circadian timekeeping *in vivo*. Hence, we initiated our investigation by delineating the neurochemical profile of DVC cells utilising RNAscope *in situ* hybridisation on tissue collected from mice at dawn (or lights-on; ZT0) and dusk (or lights-off, ZT12). Five to six DVC slices (from n=3 mice) were analysed in each time point. Previously we demonstrated using a bioluminescent reporter of the circadian clock gene protein PER2, that it is rhythmically expressed in neuron-rich and neuron-sparse areas of DVC tissue explants ^12^. However, the identity of these PER2-expressing cells remained unknown. Here, employing *Per2* as a marker for clock gene-expressing cells, we observed its high co-localisation within the DVC with both GABAergic and glutamatergic neuronal markers (identified by *Gad1* and *Slc17a6*, respectively; Fig. 1A,B). Within the AP, 85% and 83% of *Gad1*+ cells were found to co-express *Per2* (at ZT0 and ZT12, respectively). Similarly, 89% and 82% of the *Slc17a6+* population of AP cells co-localised with *Per2*. Corresponding findings were made in the NTS; 84% and 74% of putatively GABAergic, and 93% and 78% of putatively glutamatergic NTS cells co-expressed *Per2* at the two studied time points, respectively (Fig. 1C). Using single cell measurements of cell coverage by *Per2* puncta, we further showed that the level of *Per2* expression was significantly higher at ZT12 compared to ZT0 in the *Gad1*+ cells in the NTS (*p*=0.029, unpaired t-test; Fig. 1D). Similarly, we detected a high co-localisation of *Per2* with a cholinergic marker (*Chat)* within the DMV. These *Chat*+ cells showed a robust nighttime increase of the number of *Per2* puncta detected, compared to the day (*p*=0.0099, unpaired t-test; Supplementary Figure 1). This suggests that circadian timekeeping in the DVC is carried out by GABAergic, glutamatergic, and cholinergic neurons, with NTS GABAergic and DMV cholinergic cells showing a robust increase of *Per2 in vivo* from ZT0 to ZT12.

**Figure 1.**
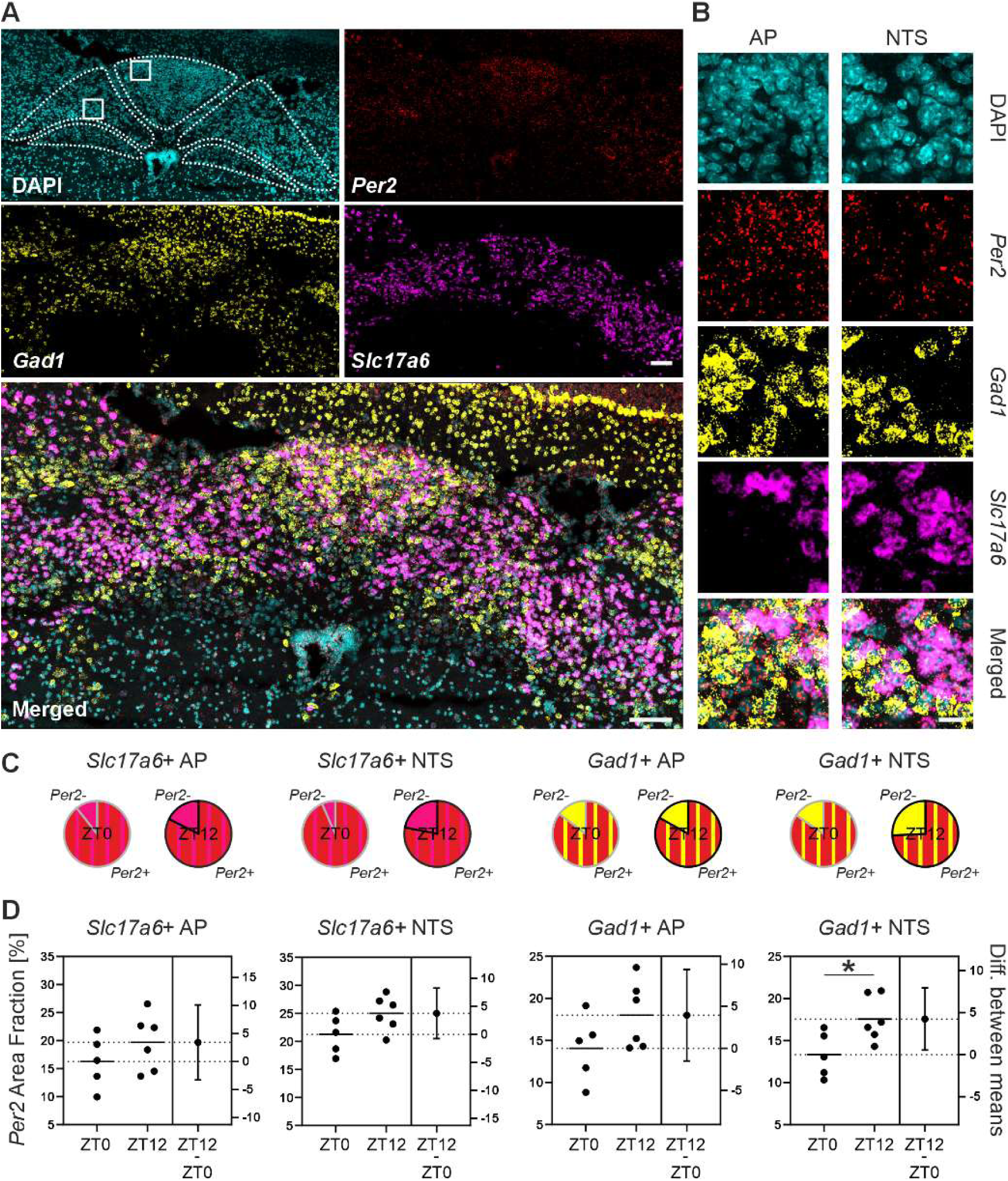
*Per2* is co-expressed with glutamatergic and GABAergic neuronal markers in the DVC. (A) Confocal microphotograph showing a representative coronal DVC section after an RNAscope *in situ* hybridisation protocol. *Gad1* and *Slc17a6* were used as markers of GABAergic and glutamatergic neurons, respectively. Both populations of DVC neurons show high co-localisation with *Per2*. White bar depicts 100 µm. (B) Zoomed images of the areas delineated by squares in *A*. White bar depicts 20 µm. (C) Pie charts depict average % of GABAergic and glutamatergic cells co-expressing *Per2.* (D) *Per2* expression level shown as area fraction (measured from single cell coverage by *Per2* and averaged within each slice), for ZT0 and ZT12. Note significant increase of *Per2* expression at the single cell level within the GABAergic population of the NTS (\**p*<0.05, unpaired t-test). All measurement were performed in n=3 animals per time point (n=1-2 slices per animal). AP – area postrema, NTS – nucleus of the solitary tract.

### Daily Rhythms in the Transcriptional Programme of the Dorsal Vagal Complex (DVC)

Clock gene proteins serve as transcriptional factors to orchestrate the timely expression of downstream genes over approximately 24h. Notably, in the SCN and other extra-SCN oscillators, the transcription of genes for receptors of neurotransmitters and neuromodulators display robust daily and circadian rhythmicity ^23–25^. Therefore, we directed our investigation towards delineating the expression of neurotransmitter receptors in tissue punches obtained from the AP and NTS at 4h intervals over a 24h period (n=5/timepoint). Utilising RT^2^ Profiler PCR Arrays, we assessed the expression levels of 84 transcripts in each sample, of which 72 and 75 were found to be reliably expressed (Ct<35) in the AP and NTS, respectively (Fig. 2A). Subsequently, by fitting sine waves to expression profiles normalised to *Gapdh* (housekeeping gene without clear circadian variation) across six daily timepoints, we identified 23 transcripts as rhythmically expressed over 24h in the AP and 32 in the NTS. A total of 15 transcripts exhibited daily rhythmicity common to both regions (Fig. 2A, Supplementary Figure 2). Intriguingly, extrapolated peak times for these rhythmic transcripts in the AP were predominantly clustered during the middle of the night, with a secondary cluster observed in the early part of the day (Fig. 2B). A similar clustering was observed for rhythmic transcripts in the NTS, albeit the peaks in the clustering were less defined (Fig. 2C).

**Figure 2.**
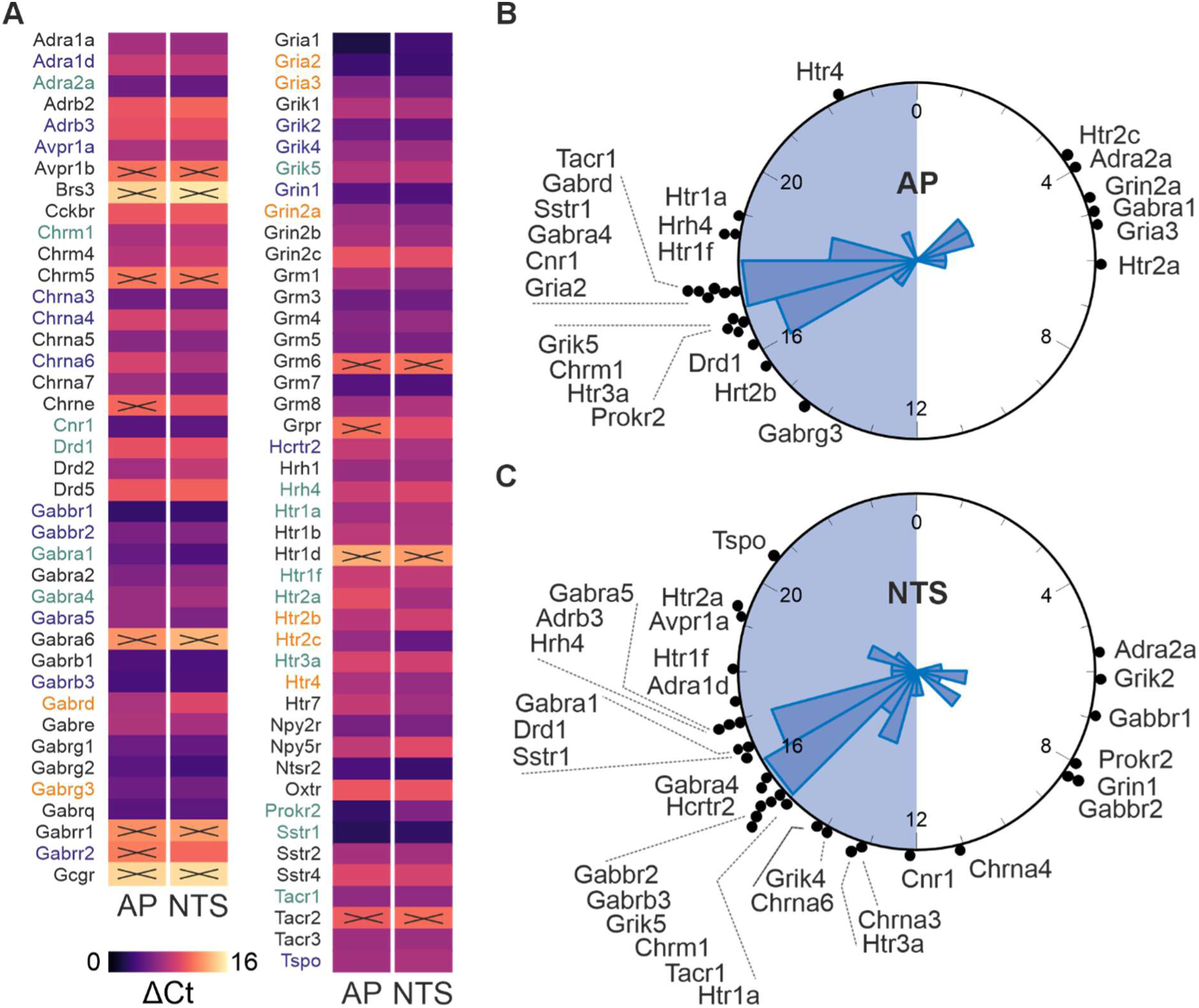
Daily regulation of the transcription of neurotransmitter receptors in the DVC. (A) Heatmap showing expression levels of investigated transcripts in the AP and NTS averaged over 24h. Low ΔCt values (Ct*^gene^*-Ct*^Gapdh^*; dark purple) depict high transcript expression, whereas high ΔCt (light yellow) – low expression. × symbol marks transcripts which did not cross the detection level (Ct*^gene^*>35). Gene names in orange followed significant rhythmic expression over 24h in the AP, in purple – in the NTS, and those in green were common for both structures. (B,C) Raleigh plots displaying acrophases for rhythmic genes in the AP and in the NTS, with circular histograms superimposed (bin=1h, bar size represents number of genes peaking at that hour). AP – area postrema, NTS – nucleus of the solitary tract. Raw data are presented in Supplementary Figure 2.

Rhythmic transcripts in the AP and NTS were further explored for the presence of putative E-boxes ^26^, and thus their potential to be directly responsive to the transcriptional activity of the molecular clock. Motif analysis assessing BMAL1 binding profile (taken from JASPAR database) revealed that 30 out of 34 rhythmic transcripts analysed contained putative E-boxes (Supplementary Figure 3). The remaining 4 genes with rhythmic transcripts (*Grik5*, *Hrt2b*, *Prok2*, and *Ssrt1*) detected with our RT^2^ Profiler PCR Arrays must be either controlled by transcription factors other than BMAL1/CLOCK or their daily oscillations are a result of rhythmic input to the DVC. Four rhythmic genes (*Adra1d*, *Adrb3*, *Grin2a*, and *Hrh4*) were excluded from the analysis, as their promoter sequences were not known or listed in the eukaryotic promoter database ^27–29^.

Previously we developed a model (based on recordings of rhythmic PER2::LUC expression in DVC tissue explants) which predicted that the circadian oscillators in the DVC would be malleable and readily reset by exogenous cues ^30^. We therefore investigated whether increased expression of neurotransmitter receptor genes in the AP and NTS is a direct consequence of feeding during the active phase of mice, commencing at the onset of the night. For this aim, we compared two cohorts of 5 animals. One group was culled immediately after undergoing a single 6h-long restriction of food availability from the late day (ZT10) to early night (ZT16). The second group experienced the same 6h food restriction followed by 2h of unlimited access to chow and thus was culled in the middle of the night (ZT18). Interestingly, this transient fasting/refeeding regimen had minimal impact on overall neurotransmitter gene expression levels in the DVC. In the AP, only 9 out of 72 transcripts tested were affected by food restriction and/or refeeding, of which just 3 exhibited daily rhythmicity in the previous experiment (Supplementary Figure 4). Similarly, in the NTS, acute changes in food availability affected 11 out of 75 transcripts, with only 5 displaying rhythmic patterns under *ad libitum* conditions (Supplementary Figure 4). These findings collectively suggest that the expression of clock genes in the AP and NTS, rather than acute change in feeding behaviour, orchestrate the majority of the phasic regulation of downstream gene expression, including those encoding neurotransmitter and neuropeptide receptors.

### Core Clock Gene Expression in the Dorsal Vagal Complex (DVC) during Time-Restricted Feeding

In contrast to acute fasting-refeeding protocols, consistent presentation of food within a predictable time window over several days induces food anticipatory activity (FAA), indicative of activation of the SCN-independent food-entrainable circadian oscillator (FEO)^9^. Although the precise anatomical location of the FEO remains elusive, it is hypothesised to be a product of a network of oscillators in central and peripheral tissues rather than a singular brain structure ^9,31^. To assess how induction of the FEO influences DVC timekeeping, we monitored the feeding, drinking, and wheel-running behaviours of mice subjected to time-restricted feeding (TRF). Initially, all mice underwent monitoring with *ad libitum* access to food under a 12:12-hour light-dark (LD) cycle as well as constant darkness (DD) to establish their daily and circadian behavioural profiles. Subsequently, following re-entrainment to LD, mice were subjected to TRF for 6-7 days, either during the initial 6 hours of the dark phase (ZT12-18, n=16) or the late portion of the light phase (ZT6-12, n=16; Fig. 3A). In both TRF groups, significant enhancement in the rhythmicity of feeding and drinking patterns were observed (Fig. 3B,C), while wheel-running rhythms exhibited reduced robustness due to fragmentation of daily locomotor activity between FAA and nocturnal wheel running (Fig. 3B,C).

**Figure 3.**
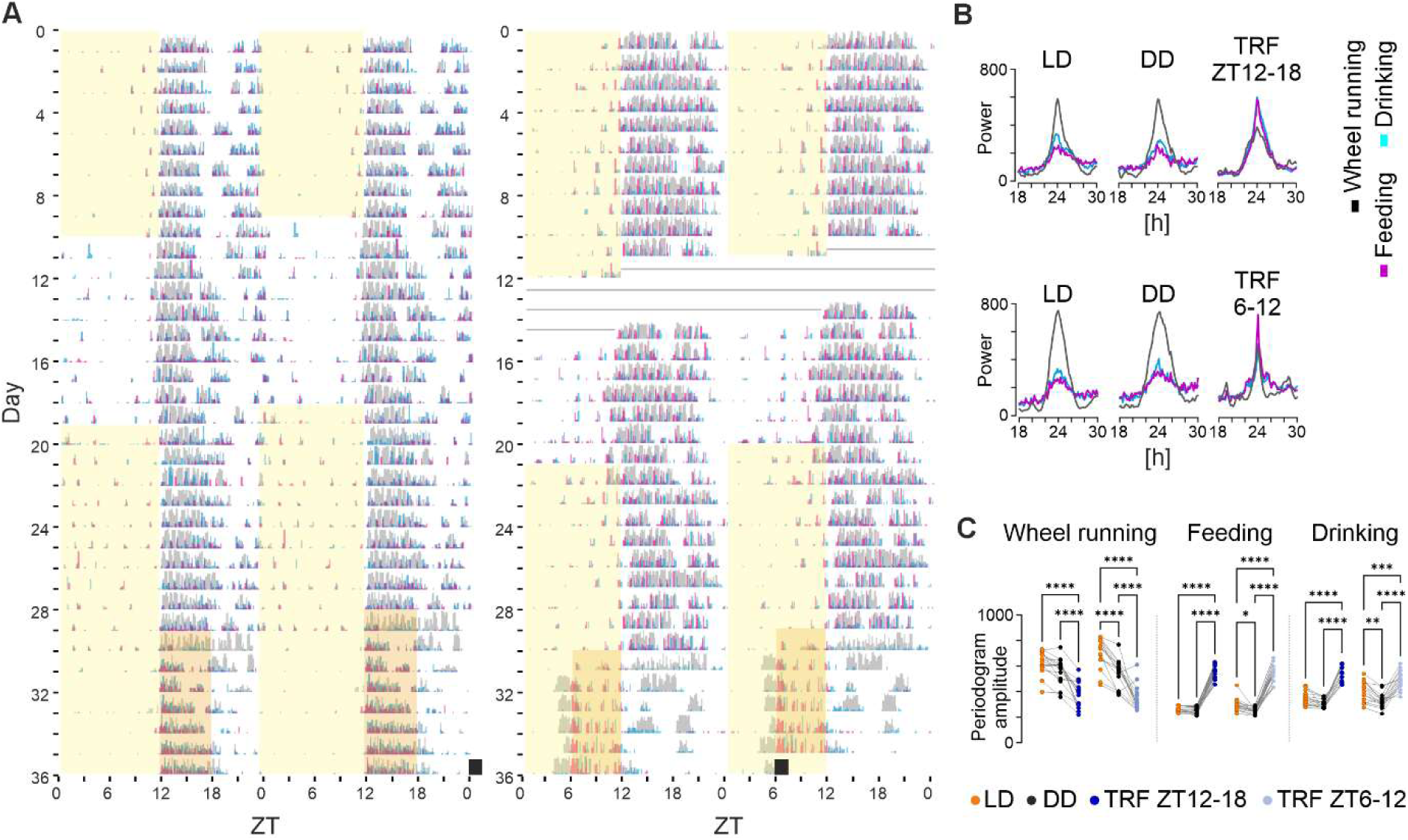
Effects of time-restricted feeding on behavioural rhythms. (A) Representative actograms showing wheel running (in grey), water (blue), and food intake (magenta). Yellow boxes depict light phase; note the persistence of behavioural rhythms in constant darkness. Straight line codes for missing data. Food was presented *ad libitum* until day 29, where 6h-long time-restricted feeding (TRF) protocol started; food presentation time is then depicted by an orange box. (B) Representative periodograms showing the robustness of ∼24h rhythms in all three behaviours under light-dark (LD), constant darkness (DD), and TRF conditions. (C) Repeated-measures comparisons showing that TRF significantly decreases robustness (periodogram amplitude) of the wheel running rhythms, while boosting feeding and drinking rhythms. \**p*<0.05, \*\**p*<0.01, \*\*\**p*<0.001, \*\*\*\**p*<0.0001, Šídák’s multiple comparisons test.

To elucidate the pattern of core clock gene expression in the DVC under these competing food and light Zeitgeber conditions, we culled mice following the same regimes of TRF at four daily timepoints (ZT0, 6, 12, and 18; n=4 each) and conducted RNAscope *in situ* hybridisation on DVC brain slices to delineate the spatiotemporal expression patterns of three core clock genes: *Bmal1*, *Per2*, and *Reverbα* (Fig. 4A). Irrespective of the timing of food presentation, all core clock genes exhibited rhythmic expression across all four regions of the DVC: the AP, NTS, DMV, and ependymal lining of the fourth ventricle (Fig. 4B). However, their phases of expression aligned with meal timing rather than light-dark cycles. Notably across all oscillators, *Reverbα* peaked approximately 6 hours prior to food presentation, while *Bmal1* peaked approximately 6 hours after food withdrawal (Fig. 4C). Interestingly, *Per2* exhibited a consistent phase across the AP, NTS, and DMV, peaking at the end of the feeding period, whereas in ependymal cells, *Per2* peaked prior to food intake in both TRF conditions (Fig. 4B,C).

**Figure 4.**
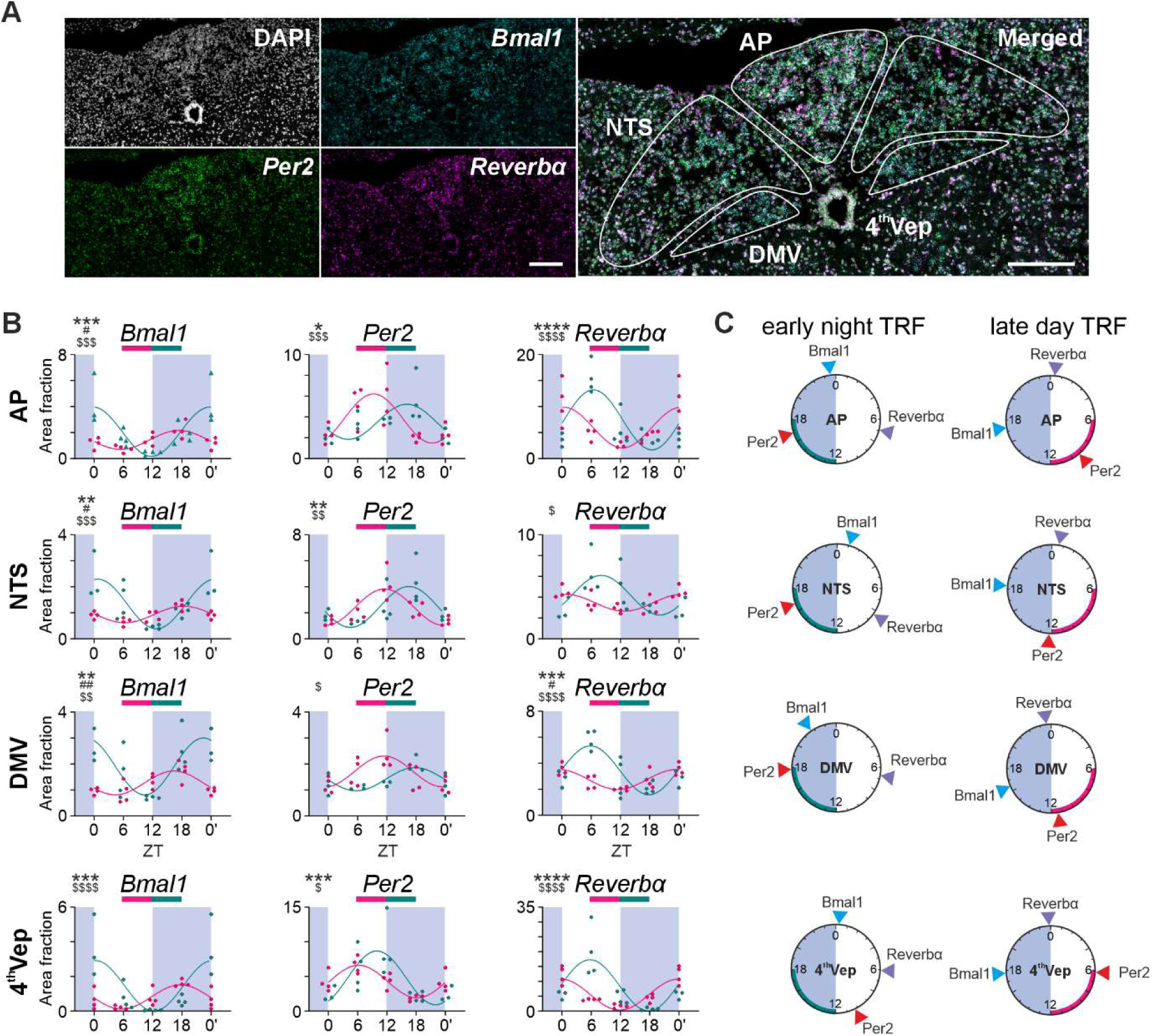
The phase of core clock genes expression in the DVC *in vivo* follows the time of food presentation. (A) Representative confocal microphotograph showing RNAscope images used for these analyses. DAPI – in grey, *Bmal1* – cyan, *Per2* – green, and *Reverbα* (magenta). Merged signal is shown on the right. White bar depicts 200 µm. (B) Sine-wave fitted area fraction measurements of rhythmic core clock gene expression in the AP, 4^th^Vep, NTS, and DMV. Magenta – late day ZT6-12 time-restricted feeding (TRF), green – early night ZT12-18 TRF. 2-way ANOVA results are depicted by * - time of day, # - feeding time, $ - interaction, where \**p*<0.05, \*\**p*<0.01, \*\*\**p*<0.001 (C) Raleigh plots displaying acrophases for core clock genes in these two TRF conditions, where the arches depict the time of food presentation. Note that peak expression of these core clock genes is better predicted by feeding time than light-dark cycle. AP – area postrema, NTS – nucleus of the solitary tract, DMV – dorsal motor nucleus of the vagus, 4^th^Vep – ependymal cell layer lining the 4^th^ ventricle/central canal.

A phase divergence between predominantly neuronal parts of the DVC and ependymal cells has been previously noted under *ex vivo* culture conditions ^12,30^. Moreover, when clock gene transcription was examined under *ad libitum* feeding conditions (n=16), the phase of *Per2* across DVC regions was synchronised, peaking around the transition from light to dark (Supplementary Figure 5). This suggests that meal timing differentially entrains the molecular clock in ependymal cells and the remaining parts of the DVC.

Next, a separate group of animals underwent early-day TRF (ZT0-6, n=16) for one week, followed by the same clock gene expression assessment with RNAscope. Surprisingly, this feeding regimen resulted in misalignment of DVC oscillators, leading to cessation of *Bmal1* rhythms in the NTS, and *Per2* rhythms in the AP and DMV (sine wave fit *p*>0.05; CircWave). With the majority of clock gene expression out of phase with food presentation compared to other TRF timings, only the remaining rhythms in the NTS reliably followed the new mealtime; *Per2* consistently peaked at the end of feeding, and *Reverbα* approximately 5 hours before food presentation (Supplementary Figure 6). With the above observation on the phase entrainment of clock gene expression by TRF, we determined a ‘food entrainment error’ (FEE, see *Methods* for description) for each gene and each DVC oscillator in order to measure the degree of entrainment to different TRF schedules. (Supplementary Figure 6). The values of FEE under late day TRF (ZT6-12) were in the range of ± 2h in all DVC substructures for all three core clock genes tested. This means that their phase can follow a 6h advance in feeding with relatively high precision. However, under an early day TRF (ZT0-6), only the acrophase of clock gene expression in the NTS showed a food entrainment error in this range, with all other structures exhibiting error in the -4h to -6h range (Supplementary Figure 6).

Taken together, these data indicate that the phase of clock gene expression in the DVC follows meal presentation time, rather than light-dark cycles. While the phase of all DVC substructures can be reliably delayed by late-day TRF, only the NTS shows enough flexibility to be advanced in phase after a one-week long early-day TRF.

### Time-Restricted Feeding Shifts Rhythms in the Transcriptional Programme in the Dorsal Vagal Complex (DVC)

Clock genes act as transcription factors, regulating the expression of numerous downstream genes to maintain proper 24-hour cycles ^5^. Our previous dataset showed that the timing of food presentation serves as a potent entraining factor for DVC molecular clock rhythms. Thus, we investigated whether TRF influences the phase of downstream rhythms in neurotransmitter receptor gene expression.

We subjected another two cohorts of mice to TRF for one week, either at the start of the light phase (TRF ZT0-6, n=16) or the dark phase (TRF ZT12-18, n=16). Seven neurotransmitter receptor genes, previously shown to exhibit robust rhythmicity under *ad libitum* feeding, were selected for analysis. Additionally, we evaluated *Per2* and *Bmal1* expression to examine the phase of the molecular clock under TRF conditions in this cohort of animals. Gene expression was analysed using NanoString nCounter technology.

Under these TRF conditions, all selected transcripts displayed significant daily rhythmicity, as determined by sine wave fitting (*p*<0.05, CircWave; Fig. 5A,B). In the early-night TRF group, most genes peaked late at night, with the exception of *Gabra1* and *Adra2a* in the AP and *Htr2c* in both the AP and NTS reaching their acrophores in the late day/early night (Fig. 5C). This bimodal acrophase distribution mirrored the pattern observed under *ad libitum* conditions (Fig. 2B,C). Notably, the phase of rhythmic neurotransmitter receptor gene expression shifted according to TRF timing (Fig. 5A,D,E). Under early-day TRF, acrophases aligned more closely with feeding times, with greater precision and clustering in the NTS compared to the AP. This was reflected in a significantly higher food entrainment error for the AP relative to the NTS (*p*=0.0223, paired t-test; Fig. 5C). As with the receptor genes, *Per2* and *Bmal1* expression also aligned with feeding schedules (Fig. 5B).

**Figure 5.**
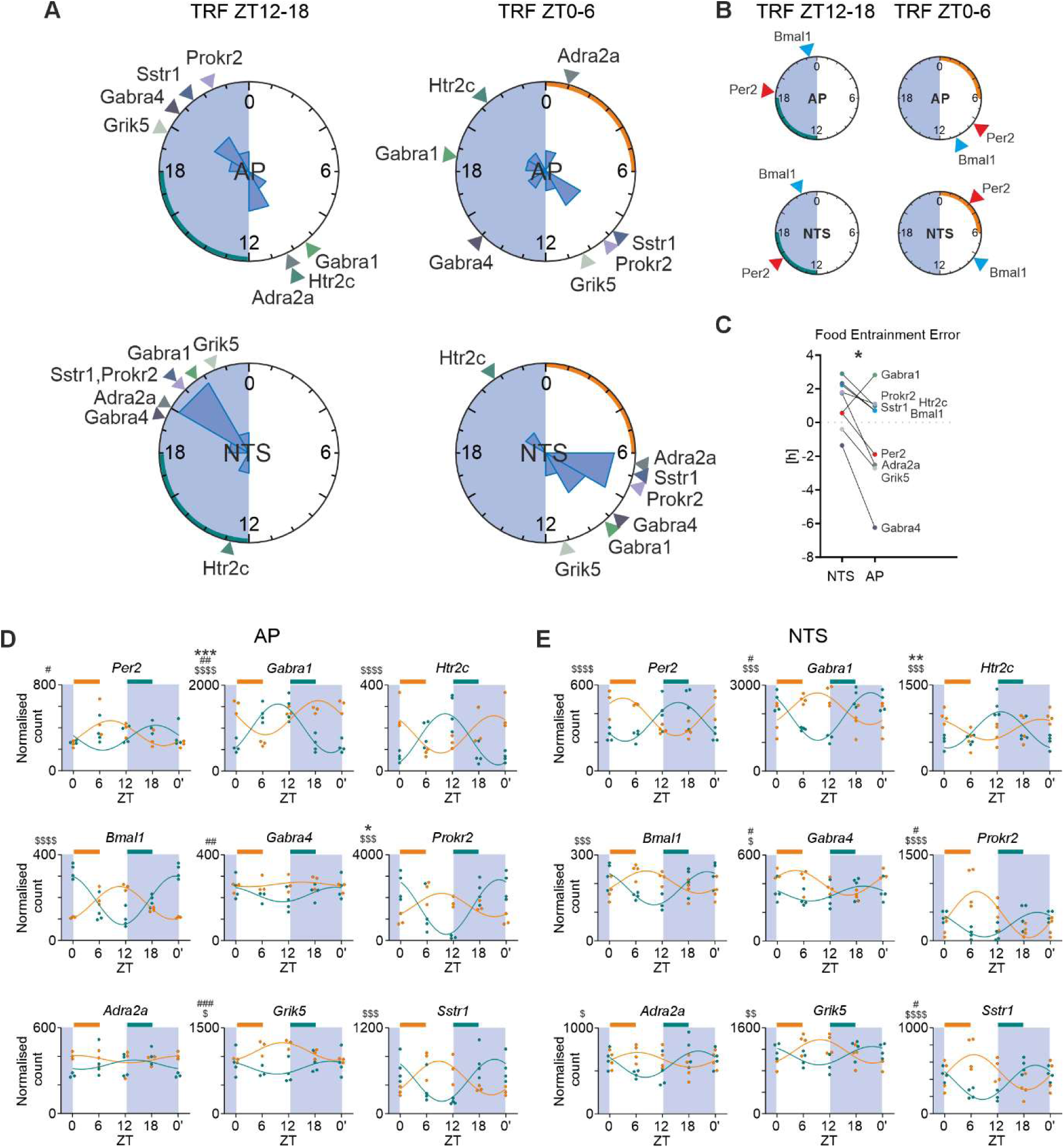
Expression of neurotransmitter receptor genes in the DVC is entrained by feeding time. (A) Raleigh plots displaying acrophases for rhythmic neurotransmitter receptor genes in the AP and in the NTS, with circular histograms (bin=2h) overlayed. Green and orange arches depict time when food was presented. (B) Corresponding Raleigh plots for core clock genes *Bmal1* and *Per2*. (C) Food entrainment error for the AP and the NTS in late day TRF condition relative to early night TRF (FEE = observed phase – projected phase); \**p*<0.05, paired t-test. (D,E) Sine-wave fitted normalised counts of rhythmic core clock transcripts in the AP and in the NTS, respectively; a result of NanoString nCounter analysis. 2-way ANOVA results are depicted by * - time of day, # - feeding time, $ - interaction, where \**p*<0.05, \*\**p*<0.01, \*\*\**p*<0.001 (and accordingly for other depicters). AP – area postrema, NTS – nucleus of the solitary tract, TRF – time-restricted feeding

Altogether, these findings demonstrate that food-entrained rhythms in molecular clock genes drive corresponding shifts in the daily oscillations of secondary genes, ensuring that neurotransmitter receptor expression in the DVC is synchronised with feeding behaviour.

## DISCUSSION

Our study reveals that the DVC demonstrates robust circadian properties *in vivo*, which are highly sensitive to meal timing. This supports the role of feeding schedules as potent Zeitgebers for extra-SCN oscillators, establishing the DVC as part of the food-entrainable clock ^8,31^. Under competing photic and non-photic Zeitgeber conditions, we show that clock gene expression in the DVC is entrained predominantly by time-restricted feeding (TRF), overriding light-dark cycles. The key finding of this work is that this food entrainment affects the timely transcription of neurotransmitter receptor genes, aligning the DVC transcriptome with feeding behaviour.

The DVC is a critical hub for metabolic, cardiovascular, and ingestive processes essential for survival. Its medial part containing the AP contributes specifically to meal termination and serves as both a source and a primary target of vagal circuitry connecting the gastrointestinal tract and the brain. Additionally, neurons within the DVC respond to peri-prandial hormonal cues, further underscoring its role in the gut-brain axis ^13,14,32,33^. Our findings reveal that the molecular clock is expressed in multiple neuronal populations within the DVC, including GABAergic and glutamatergic cells in the AP and NTS, and cholinergic neurons in the DMV. These findings extend our previous observations ^12^, demonstrating widespread circadian timekeeping across the DVC that parallels the SCN, where diverse neuronal populations synchronise molecular rhythms to regulate physiological outputs ^34^

Our prior *ex vivo* and *in vivo* work, as well as computational modelling studies ^12,18,19,30,35,36^ have established the intrinsic circadian properties of the DVC. However, the environmental cues that entrain the DVC remained unclear until now. In the SCN, synchronised rhythms in molecular clock expression, which maintain daily outputs such as neurophysiological activity and neurohormone release, are entrained predominantly by light ^6,37^. In contrast, the DVC lacks direct retinal innervation^38,39^ and it has a well-established role in controlling ingestive behaviour ^13,14^.

Accumulating research has shown that long-term high-fat diet (HFD) disrupts circadian oscillators ^40,41^, including molecular clock rhythms in the DVC ^20^. Moreover, it is well-documented that feeding patterns are altered by consumption of HFD via dysfunction of the molecular clock as well as an increased hedonic drive to eat highly palatable food throughout the 24h cycle ^42^. Our own studies corroborate these findings, showing that even short-term HFD alters rhythmic properties in the DVC *ex vivo* while disrupting feeding patterns, leading to food intake during the behaviourally quiescent light phase ^18,19^. Here we show that meal timing is crucial for the phase of clock gene expression in the DVC, which suggests that changes in daily patterns of feeding evoked by HFD, rather than diet composition, may contribute to the detrimental effects of HFD on the DVC circadian timekeeping. Therefore, the potential protective effects of TRF during the behaviourally active phase on the rhythmicity of the DVC under HFD conditions warrant further investigation.

Our findings further support the inclusion of the DVC within the food-entrainable oscillator (FEO) network. The alignment of DVC clock gene expression to TRF is consistent with patterns observed in hypothalamic feeding centres including the arcuate nucleus and dorsomedial hypothalamus ^43–45^. While the SCN-independent nature of FAA has been well-established ^9^, the specific role of the DVC within this network remains to be fully elucidated. It is unlikely that the AP is critical for the expression of FAA, as a previous study demonstrated that AP lesions do not abolish this behaviour ^46^, reinforcing the idea that no single brain structure constitutes the entire FEO ^11^. However, further studies using genetic models are needed to investigate possible involvement of other DVC nuclei (particularly the NTS) in the generation of FAA. To date, our findings position the DVC as a flexible node in the FEO network, capable of adapting its transcriptional program to predict meal availability.

Interestingly, the timing of TRF within the 24-hour cycle affects the degree of DVC entrainment. A late-day TRF schedule (phase advance relative to the active phase) produced more reliable shifts in DVC rhythms compared to early-day TRF (phase delay). This aligns with behavioural evidence suggesting that circadian rhythms are more readily advanced by non-photic cues and delayed by photic stimuli ^47^. Although a week of TRF is enough to synchronise peripheral clocks to a reversed feeding time ^48^, longer TRF protocols may enhance entrainment to early-day feeding schedules. Among DVC components, the NTS demonstrated the most precise alignment of clock gene expression under both TRF conditions, suggesting a role for vagal afferent signalling in entraining the DVC circadian timekeeping. Given that gut-derived vagal innervation preferentially targets the NTS ^14,49,50^, it is plausible that the vagus nerve functions as a key pathway for feeding-driven entrainment, analogous to the role of retinal input in the SCN. Future studies are needed to investigate this hypothesis.

Crucially, these food-entrained molecular clock rhythms were shown to orchestrate the rhythmic transcriptional repertoire of the DVC. Under *ad libitum* feeding, we observed daily oscillations in the expression of multiple neurotransmitter receptor genes. Similar temporal clustering or synchronisation of gene expression has been reported in other clock gene-expressing peripheral tissues, such as the heart and liver, where the circadian clock governs tissue-specific functions ^51^. Our findings also mirror those in the SCN, where the molecular clock’s role in regulating downstream gene expression is well established ^4,5^. Importantly, the phase of these gene expression rhythms in the DVC shifted in accordance with clock gene entrainment to altered meal timing. This includes transcripts encoding neurotransmitter receptors with documented roles in the control of feeding (such as *Sstr1, Hrt2c,* and *Cnr1*)^52–54^ and gastric mobility (*Gabra1* or *Gabra4*)^55,56^. This highlights the functional output of the DVC’s circadian machinery, shaping the daily transcriptional landscape of its constituent cells to align with environmental feeding cues.

Our findings have important implications for understanding the circadian plasticity of the DVC and its role in metabolic health. Feeding schedules that entrain DVC molecular clocks could optimise the timing of neurotransmitter receptor expression, potentially enhancing the efficacy of pharmacological interventions targeting DVC circuits. Pharmacological agents that target neurotransmitter receptors in the DVC including CNR1, HTR2c, and GLP1R are already in clinical use or under investigation for obesity and diabetes treatment ^15,17,57^. Incorporating meal timing into chronopharmacological strategies could synergise with these treatments, utilising the DVC’s circadian rhythmicity to improve therapeutic outcomes.

In conclusion, our study establishes the DVC as an integral node in the FEO network, capable of integrating feeding cues to modulate circadian gene expression. By demonstrating the entrainment of DVC molecular clocks by meal timing, we provide new insights into the neural mechanisms underlying food entrainment and lay the groundwork for circadian-based interventions in metabolic disorders.

## METHODS

### Animals

All experiments described in this article were conducted using adult (>8 weeks old) mice of both sexes, on a C57BL/6J genetic background locally bred at the Animal Services Unit at the University of Bristol. Mice were housed at 20–22°C with around 40% humidity, provided with *ad libitum* access to food (unless otherwise specified) and water before and throughout the experiments, and maintained on a 12:12-hour light-dark (LD) cycle in breeding rooms. Animals were also kept under 12:12-hour LD conditions during experiments unless otherwise stated. All experimental procedures were conducted in accordance with the UK Animals (Scientific Procedures) Act of 1986 and project-specific licences. All measures were taken to refine procedures and minimise animal suffering.

### Fluorescent in situ hybridisation (RNAscope)

#### Tissue Preparation

Twelve mice were deeply anaesthetised with an overdose of pentobarbital sodium (200 mg/ml, i.p.) at ZT0 (n=6) and ZT12 (n=6), and subsequently decapitated. Brains were promptly removed from the skulls, flash-frozen in Cryomatrix (Epredia, UK) over dry ice, and stored at -80°C. This set of tissue was utilised to characterise clock gene expression in GABAergic vs glutamatergic neuronal populations in the DVC (*Experiment 1*, n=3 at ZT0 and n=3 at ZT12), and to investigate the co-localisation of clock gene expression with *Chat* (*Experiment 2*, n=3 at ZT0 and n=3 at ZT12). Two slices were analysed from each mouse.

Another set of RNAscope hybridisation (*Experiment 3*) was performed to elucidate rhythms in clock gene expression in the DVC of mice undergoing 6h-long time-restricted feeding (TRF) for 7 consecutive days. 64 mice were divided into four feeding groups (*ad libitum*, TRF ZT0-6, TRF ZT6-12, TRF ZT12-18) and humanely killed in 6-hour intervals over 24 hours (ZT0, 6, 12, and 18; n=4 animals per timepoint per diet). Two to three slices were analysed from each animal and averaged within each mouse.

All brains were sectioned into 16 μm thick coronal slices at -20°C using a cryostat (Leica CM1860 UV). Slices were thaw-mounted on Superfrost-Plus slides (Thermofisher, USA), and stored at -80°C until the day of the protocol. Then, slices were thawed, fixed in 4% paraformaldehyde (PFA) solution in 0.1M phosphate buffered saline (PBS) for 15 minutes at room temperature (RT), rinsed twice in fresh PBS, and dehydrated in increasing ethanol concentrations (50%, 70%, 100%, and 100%). Slices were stored at -20°C in the second 100% ethanol solution overnight.

#### RNAscope Protocol and Imaging

First, slides were air dried, and each slice was outlined with a hydrophobic barrier pen. Then, slices were processed according to RNAscope multiplex v2 in situ hybridisation protocol (Advanced Cell Diagnostics—ACD, USA). In brief, slices were incubated at RT with hydrogen peroxide for 10 minutes and with protease IV for a further 12 minutes. Slices were then incubated at 40°C for 2 hours with a set of probes depending on the Experiment: (*1*) *Per2, Gad1*, and *Slc17a6,* (*2*) *Per2* and *Chat,* (*3*) *Arntl, Per2*, and *Nr1d1*. Next, the signal was amplified in a three-step protocol. Finally, up to three fluorophores (Opal dyes: 520, 570, and 690; Akoya Biosciences, MA, USA) were tagged via a horseradish peroxidase reaction. Slides were air dried, and cover slipped with Fluoroshield containing DAPI (Sigma, Germany).

#### Imaging, Analysis, and Statistics

The area containing the DVC was scanned at 40 × magnification with the confocal scanning system (Leica SP8, Leica Microsystems, Germany), and images were captured using Leica Application Suite X (LasX, Leica Microsystems). Typically, 18 tile images were captured per DVC, with each tile consisting of 6-8 images in a z-stack (z-step = 1 µm).

Images were further analysed in FIJI (ImageJ) using a custom-made MIA plugin. Cell counting and co-expression analysis were performed at a single cell level for *Experiment 1* and *2*, whereas *Experiment 3* was analysed by drawing polygon regions of interest (ROIs) delineating the whole DVC subdivisions (the AP, NTS, DMV, and 4thVep).

Numerical results were statistically analysed using RStudio v4.3.0 and GraphPad Prism 10 (GraphPad Software, MA, USA), and rhythmicity in clock gene expression (*Experiment 3*) was assessed using CircWave v1.4 (Dr. Roelof Hut, http://www.euclock.org/).

Relation of CircWave-determined acrophases of gene expression to feeding time were measured as a food entrainment error (FEE). FEE was calculated as a difference between the experimentally observed and the expected acrophase, relative to early night TRF. For example, a FEE would equal 0, if the peak gene expression was advanced by exactly 6h following a 6h phase advance of feeding from early night TRF (ZT12-18) to late day TRF (ZT6-12).

### RT^2^ Profiler PCR Arrays

#### Tissue Preparation

Thirty mice were fed *ad libitum* and culled at six daily timepoints with 4-hour intervals over 24 hours (n=5 per group) by an overdose of pentobarbital sodium (i.p.; *Experiment A*). An additional group of ten animals were acutely food restricted at ZT10, with a subgroup of five culled after 6 hours (at ZT16; fasted), and the remaining five regaining access to food for 2 hours before being humanely killed at ZT18 (re-fed; *Experiment B*). All 40 brains were extracted from the skull in ice-cold Hanks’ balanced salt solution (HBSS, Sigma, UK). Hindbrains were cut using a vibroslicer (Campden Instruments, UK) into 250 µm-thick slices containing the DVC, from which the AP and the bilateral NTS were collected using a sample corer (Fine Science Tools, Germany; ID: 0.5 mm). 80 samples of tissue were then flash-frozen over dry ice and stored at -80°C.

#### RNA Extraction and RT-qPCR

Tissue was processed for RNA extraction using the ReliaPrep RNA Tissue Miniprep System (Promega, USA). Extracted RNA in RNase-free water was stored at -80°C until reverse transcription with the High-Capacity RNA-to-cDNA Kit (Applied Biosystems, USA). Subsequently, cDNA was stored at -20°C. The RT-qPCR reaction was performed using RT^2^ Profiler PCR Arrays (Qiagen, USA) in the Mouse Neurotransmitter Receptors configuration (GeneGlobe ID: PAMM-060Z) – a preloaded PCR plate with qPCR primers targeting 84 genes coding a range of neurotransmitter receptors. Thermal cycling and data collection were performed using QuantStudio3 (Invitrogen).

#### Analysis and Statistics

Results were then analysed according to the Livak method (2^−ΔΔCT^) with *Gapdh* as the reference gene and presented as relative target gene expression (RQ), where RQ = 1 indicates the mean expression of a gene of interest at ZT0 (*Experiment A*) or at *ad libitum* fed ZT16 (*Experiment B*). Genes were classified as not expressed if their mean Ct throughout six daily timepoints >35. RQ values were further analysed in GraphPad Prism 10 to evaluate significant variability of expression between groups, and sine-wave fitted with CircWave v1.4 to assess daily rhythmicity in gene expression.

Motif analysis was conducted using the Biopython package (v1.78) in Python (v3.9.9) ^58^. Instances of the BMAL1 binding profile were assessed in the promoter and 5’ untranslated sequence of the genes, 1kb either side of the transcription start side, with a false negative rate of 0.1. The BMAL1 binding profile used was from the JASPAR database (9th release; matrix ID MA0603.1) and the sequences were collected from the eukaryotic promoter database ^27–29^.

### Feeding, Drinking, and Wheel Running Assessment

#### Behavioural Protocol and Time-Restricted Feeding

32 animals were single-housed in running wheel-equipped cages with a precision balance to monitor drinking and feeding activity (TSE Systems, Germany). Wheel running, drinking, and feeding activities were recorded using PhenoMaster software (TSE Systems) following a similar experimental set-up as reported before ^59^. All mice were first monitored for at least 10 days in a 12:12-hour LD cycle under *ad libitum* feeding conditions, and then their behavioural activities were recorded in constant darkness (DD) for a further 10 days. After this period, the 12:12-hour LD was re-established for the rest of the experiment. Following another 10 day-long epoch, mice were divided into two time-restricted feeding (TRF) cohorts (n=16 each) with different times of food presentation. The first group had unlimited access to food for the last six hours of the light phase (TRF ZT6-12), whereas the second cohort was fed during the first six hours of the night (TRF ZT12-18). Animals underwent TRF for 7 days before being culled at four daily timepoints (ZT0, 6, 12, and 18). Water was provided *ad libitum* throughout the entirety of all protocols. Brain tissue was further used for RNAscope *Experiment 3* (see: Fluorescent in situ hybridisation (RNAscope)).

#### Behavioural Analysis and Statistics

The circadian period and power (the amplitude of the rhythm) of the behavioural data were assessed with a chi-square periodogram using Actimetrics Clocklab software (v6.1.10; Lafayette Instrument Company, Lafayette, IN, USA). Outliers in the feeding and drinking data were removed with an exclusion criterion of above 12SD from the mean average of an active period using R (R v4.4.1). Statistical analysis of data was performed in GraphPad Prism 10.

### NanoString nCounter

#### Tissue Preparation and RNA extraction

Another group of 32 mice were single housed with *ad libitum* access to food and water, under a 12:12h LD cycle. Then, animals were divided into two cohorts (n=16 each). The first was subjected to TRF from ZT0 to ZT6 (early day), whereas the second group was fed between ZT12 and ZT18 (early night). Following 6-7 days of TRF, animals were culled at four daily timepoints in 6-hour intervals (ZT0, 6, 12, and 18). Tissue was processed in the same way as for RT-qPCR, and RNA extracted from 32 AP and 32 NTS tissue punches was stored at -80°C.

#### Probe design, sample processing, and analysis

For each reaction, we used a combination of 11 probes: seven targeting neurotransmitter receptors (*Htr2c, Grik5, Gabra4, Gabra1, Adra2a, Prokr2, Sstr1*), two - molecular clock components (*Arntl* and *Per2*) and two - housekeeping genes (*Actb* and *Gapdh*). Oligonucleotides were purchased separately from Integrated DNA Technologies (IDT, USA). Samples were processed by the Genomics Core Facility at the University of Bristol according to nCounter Elements TagSets protocol provided by NanoString. Data were analysed using nSolver 4.0 software (NanoString). Raw counts were first thresholded with a geometric mean of the negative control counts, and then normalised to positive controls and housekeeping genes. Data were presented as relative target gene expression (RQ), where RQ = 1 indicates the mean expression of a gene of interest at ZT0. RQ values were further plotted and statistically analysed in GraphPad Prism 10, and sine-wave fitted with CircWave v1.4.

## Acknowledgements

This study was financially supported by Sir Henry Wellcome Postdoctoral Fellowship (Wellcome Trust, UK; 224116/Z/21/Z) awarded to LC. TH was supported by a BBSRC grant to HDP (BB/R019223/1), CM and TCK are supported by BBSRC SWBio Doctoral Training Program studentships (BB/T008741/1).

---

**Supplementary Figure 1.**
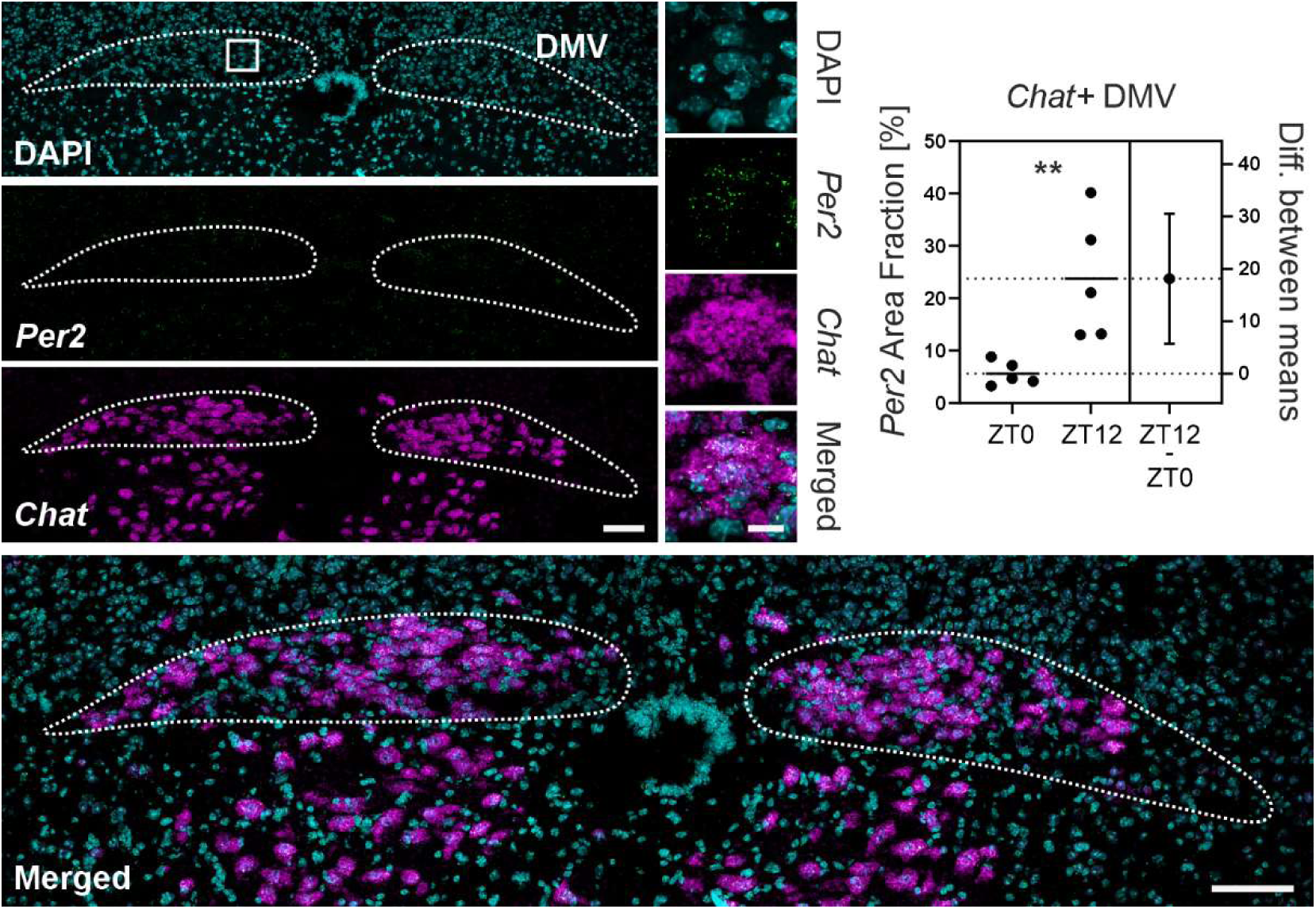
*Per2* is expressed by cholinergic DMV neurons. RNAscope confocal microphotograph showing the co-localisation of *Per2* (in green) with *Chat* (a cholinergic marker; in magenta). Additionally, a robust day to night difference in *Per2* expression within the *Chat*+ cells is shown (\*\**p*<0.01, unpaired t-test). White bar depicts 100 µm whereas in the zoomed images (magnifying the area depicted by the square) on the right –20 µm.

**Supplementary Figure 2.**
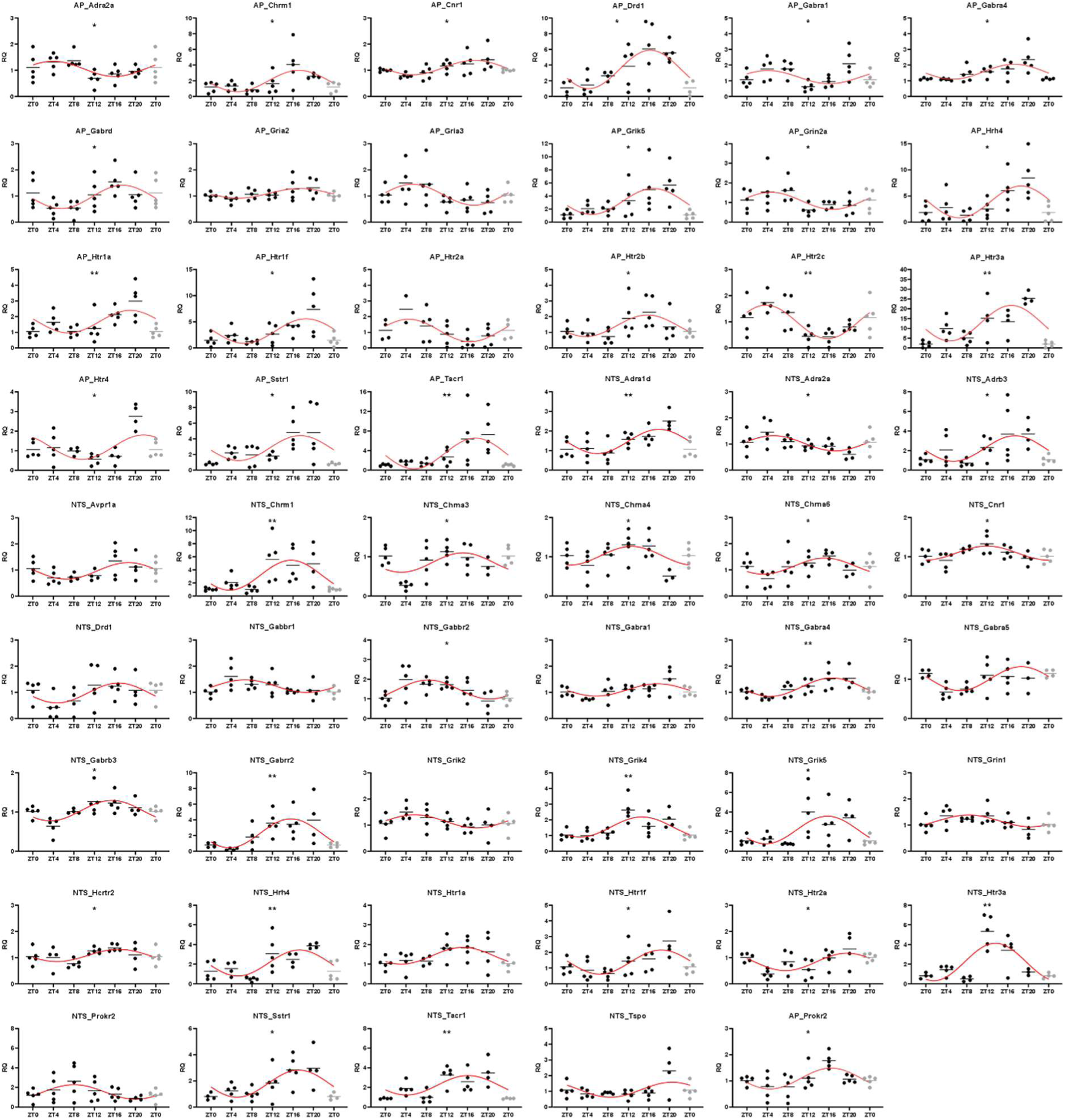
Rhythmic expression of neurotransmitter receptor genes in the AP and NTS. Sine-wave fitted normalised counts of rhythmic neurotransmitter receptor transcripts in the AP and in the NTS. Data were collected using RT qPCR Profiler Arrays. RQ – gene expression relative to ZT0. \**p*<0.05, \*\**p*<0.01, Kruskal-Wallis test.

**Supplementary Figure 3.**
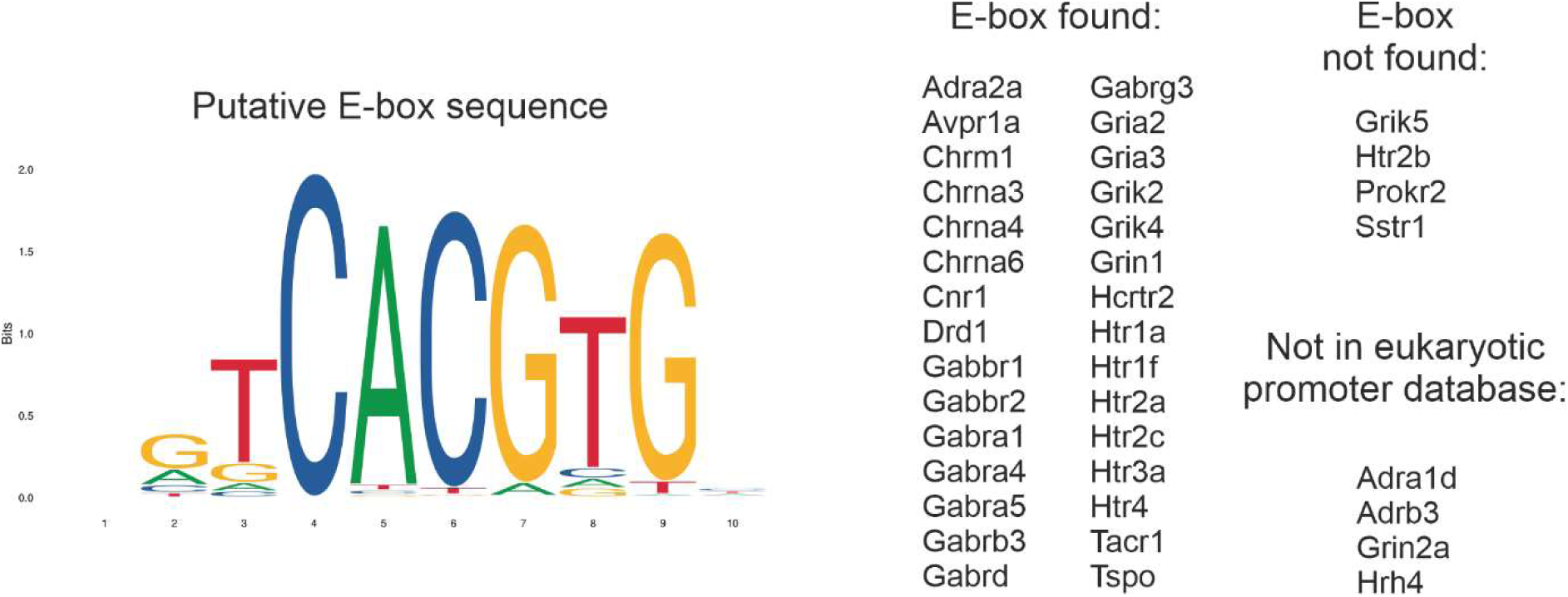
Enrichment analysis of JASPAR transcription factor binding sites (TFBSs). (*left*) TFBS sequence motif for a putative E-box sequence is illustrated with the height of each base indicative of the probability of their presence at the designated position. (*right*) A list of genes showing results of the putative E-box sequence search.

**Supplementary Figure 4.**
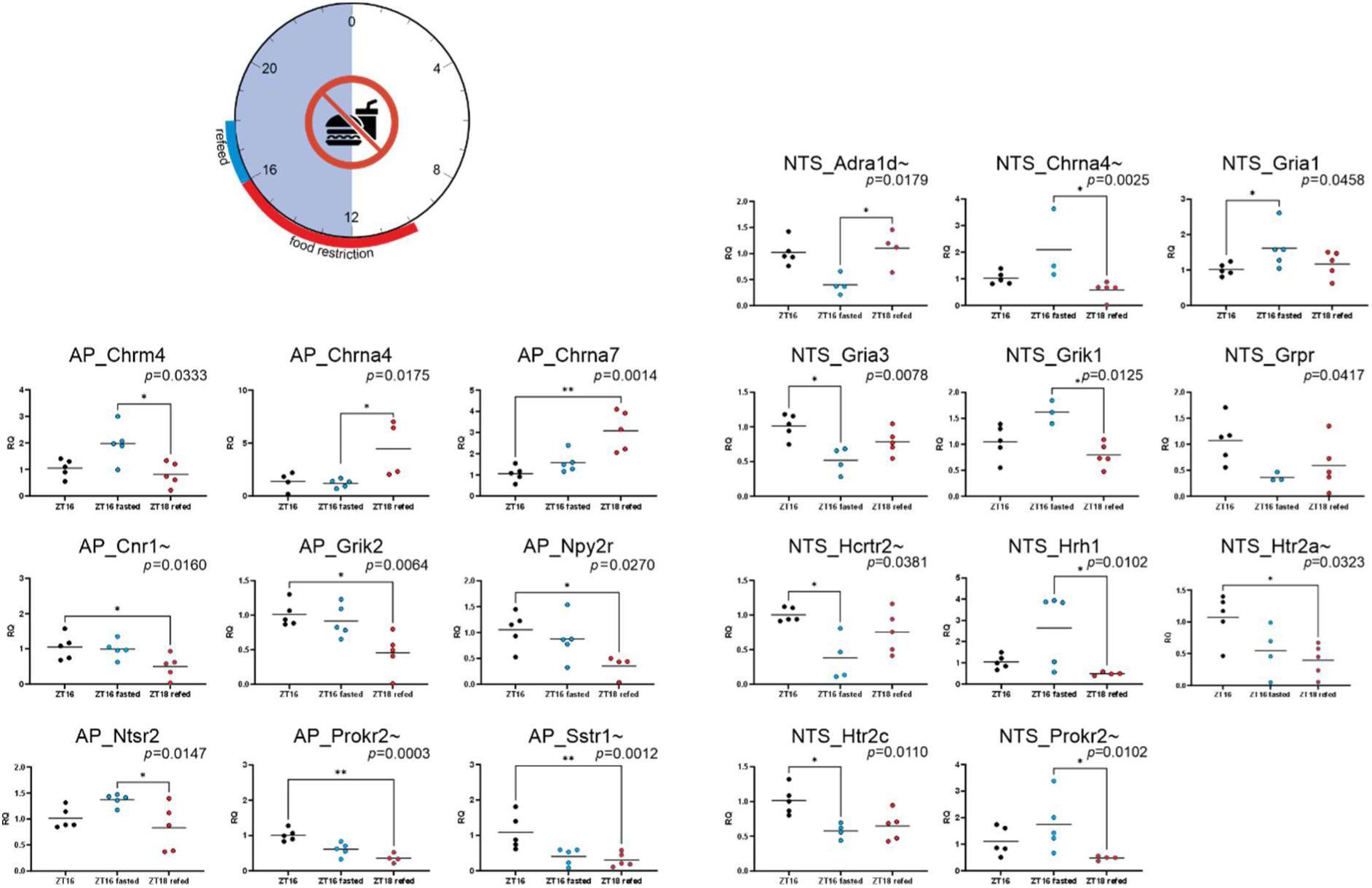
Effects of a one-off food restriction on the transcription of neurotransmitter receptor genes in the DVC. The cartoon shows an experimental timeline – animals were food restricted between ZT10 and ZT16, and some of them were re-fed 2h prior to the cull. Scatter plots show the result of RT qPCR Profiler Array comparison of relative gene expression (RQ, normalised to ZT16) between ZT16 with (black)and without (blue) access to food, and the refeeding condition (red). Kruskal-Walis test results are presented as *p* values in the graphs, whereas * show the result of Dunn’s multiple comparisons test. Only genes significantly changed by acute feeding conditions (as depicted by main ANOVA result) are presented. ∼ next to the gene name depicts that this transcript was deemed to be rhythmic under *ad libitum* conditions, when evaluated over 24h (see Fig. 2).

**Supplementary Figure 5.**
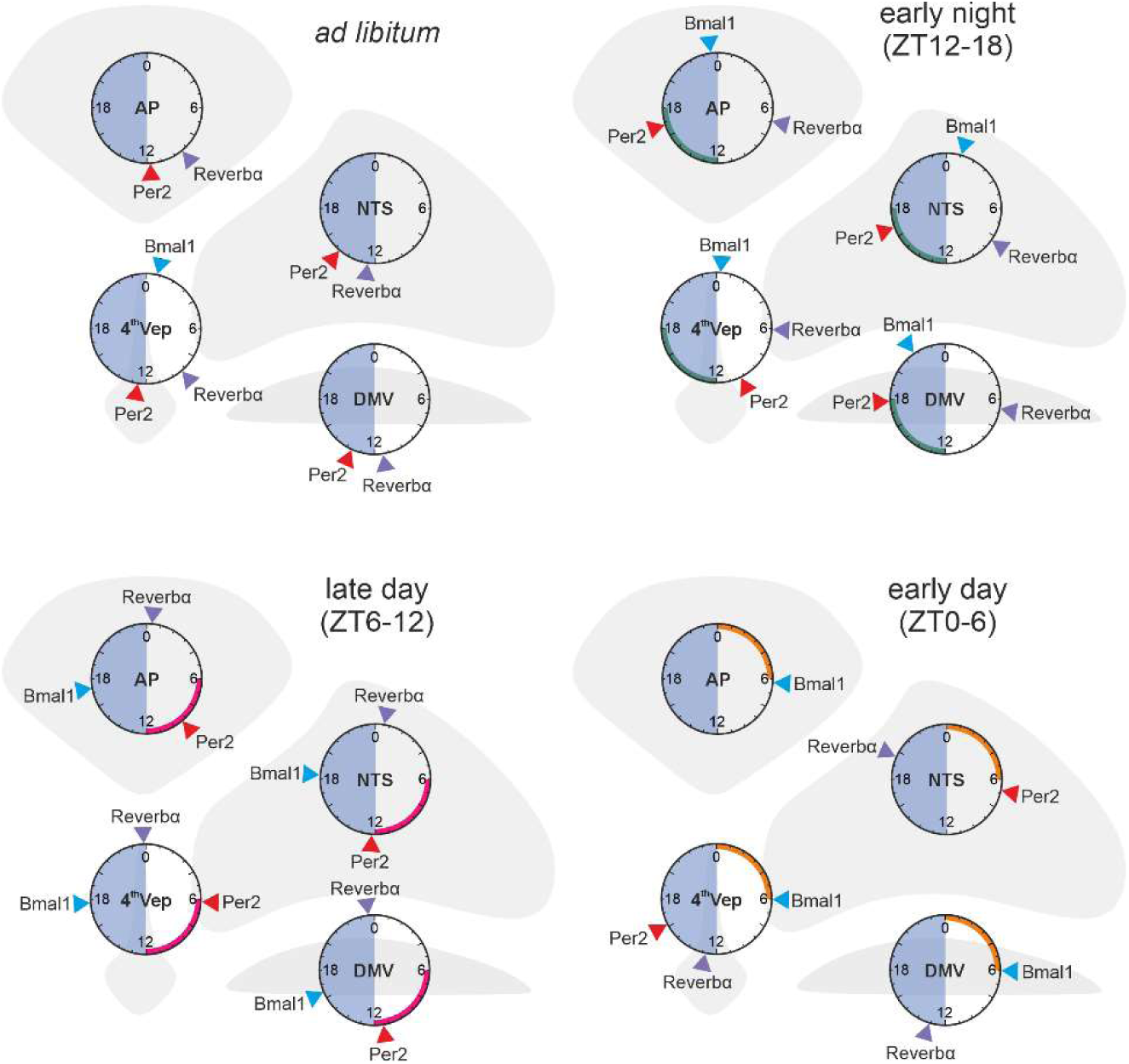
Summary of acrophases of *Bmal1*, *Per2*, and *Reverbα* expression in four parts of the DVC under different food presentation times. Raleigh plots display peak expression for core clock genes under *ad libitum* feeding (*top left*) in three time-restricted feeding (TRF) conditions. Arches within the Raleigh plots depict TRF time: green – early night ZT12-18, magenta – late day ZT6-12, and orange – early day ZT0-6. Note that genes which were not significantly rhythmic (CircWave *p*>0.05 for a sine wave fit) were not plotted. Data were collected using area fraction measurements on images obtained with RNAscope *in situ* hybridisation. AP – area postrema, NTS – nucleus of the solitary tract, DMV – dorsal motor nucleus of the vagus, 4^th^Vep – ependymal cell layer lining the 4^th^ ventricle/central canal.

**Supplementary Figure 6.**
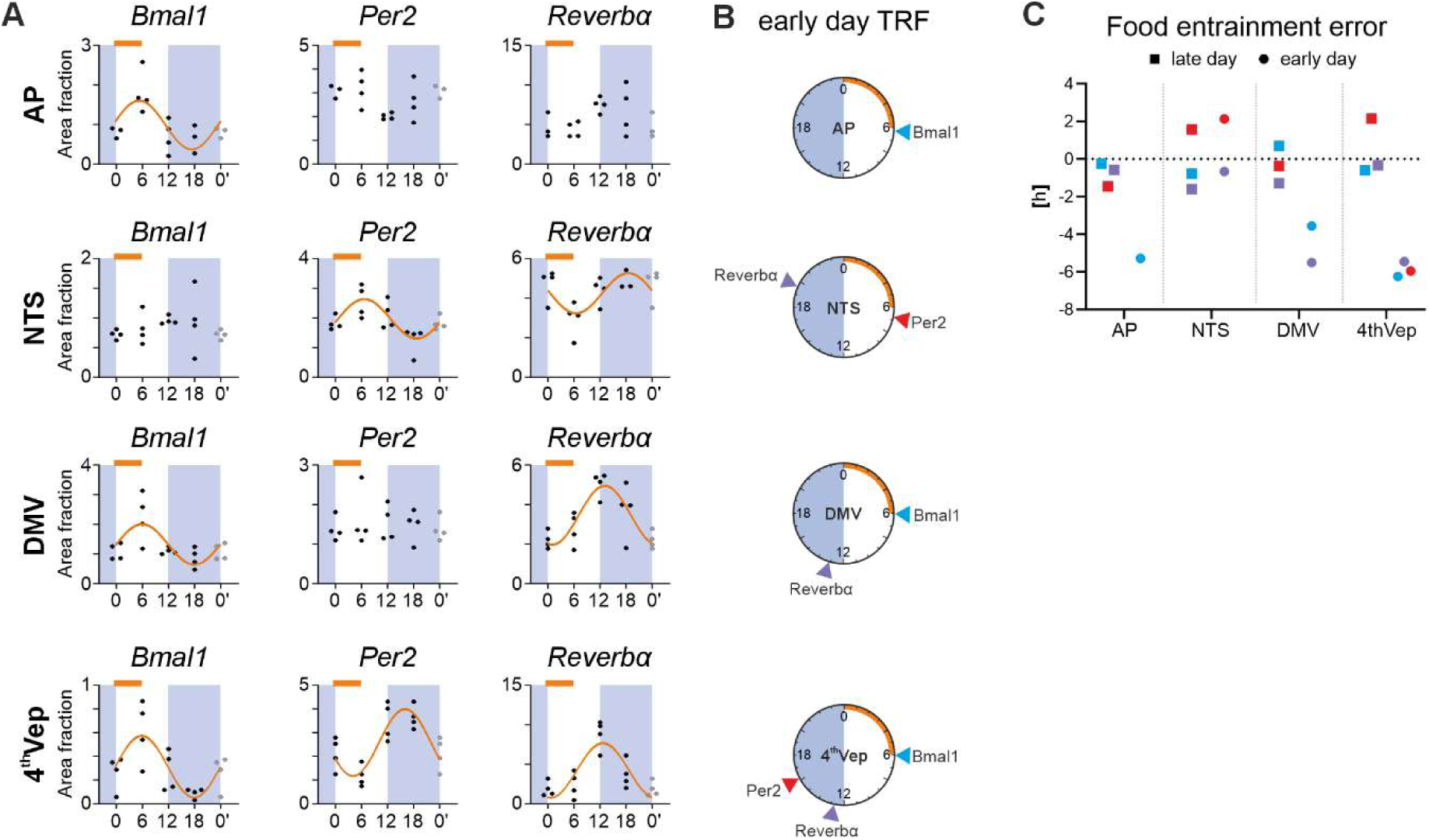
Core clock gene expression in the DVC over 24h under early day time-restricted feeding (TRF). (A) Orange bar depicts the TRF epoch, whereas significant fit of an orange sine wave (CircWave *p*>0.05) shows rhythmicity of a transcript across 24h. Data were collected using area fraction measurements on images obtained with RNAscope *in situ* hybridisation. (B) Raleigh plots displaying acrophases for core clock genes in early day TRF, where the orange arches depict the time of food presentation. Note that genes which were not significantly rhythmic were not plotted (C) Food entrainment error relative to early night TRF (FEE = observed phase – projected phase) for *Bmal1* (blue), *Per2* (red), and *Reverbα* (purple) for late day (squares) and early day TRF (circles). (E) AP – area postrema, NTS – nucleus of the solitary tract, DMV – dorsal motor nucleus of the vagus, 4^th^Vep – ependymal cell layer lining the 4^th^ ventricle/central canal.

## Notes

### Competing Interest Statement

The authors have declared no competing interest.

