## Supplementary figures and images for "Food-entrainment of circadian timekeeping in the dorsal vagal complex"

### Supplemental Figure 1

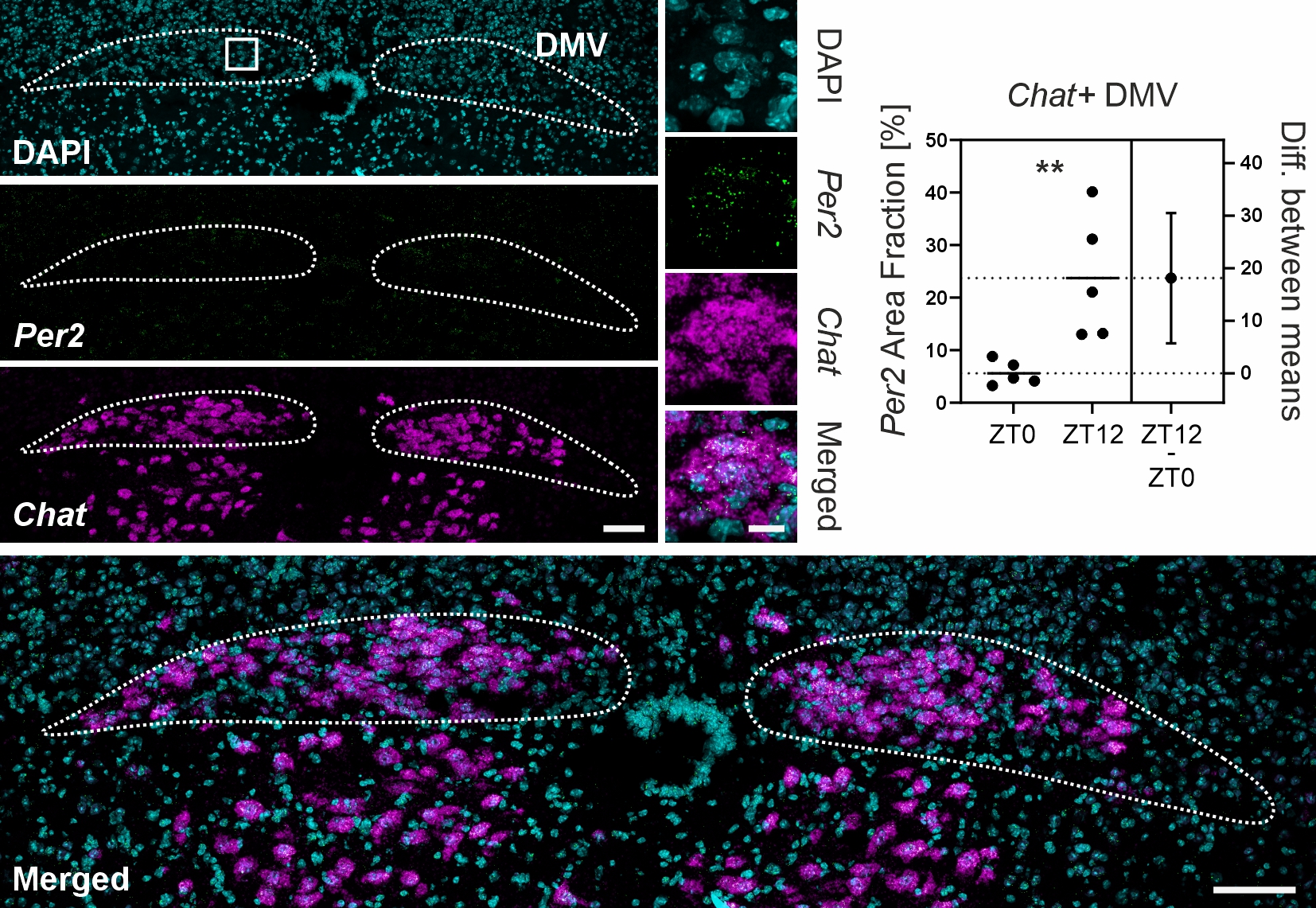

### Supplemental Figure 2

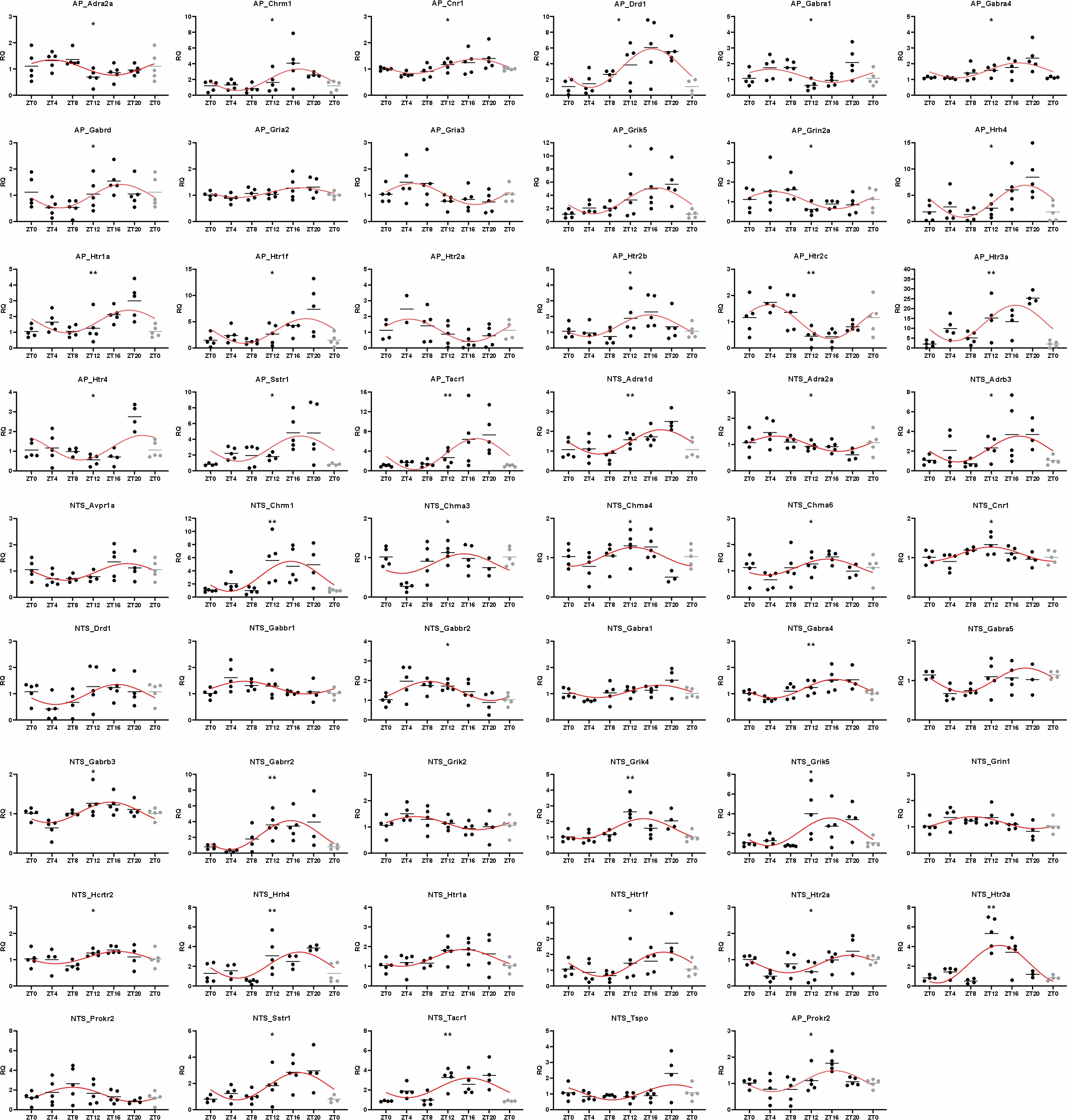

### Supplemental Figure 3

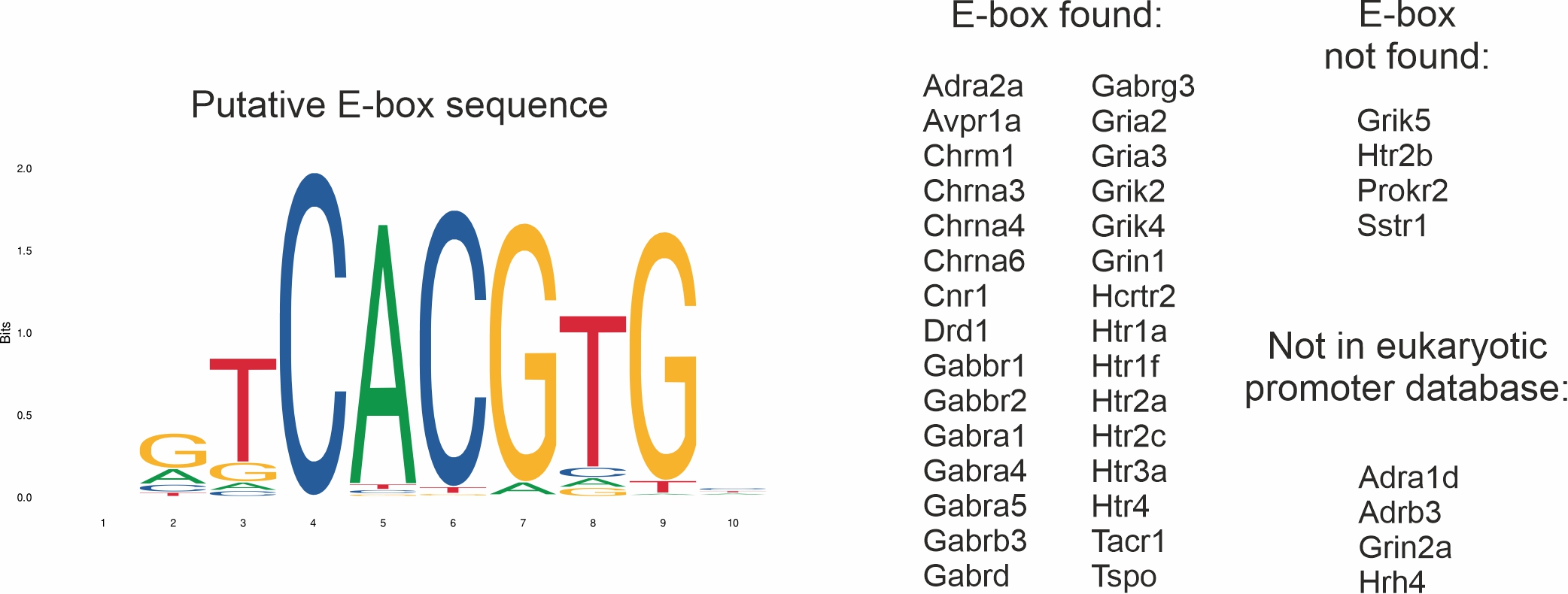

### Supplemental Figure 4

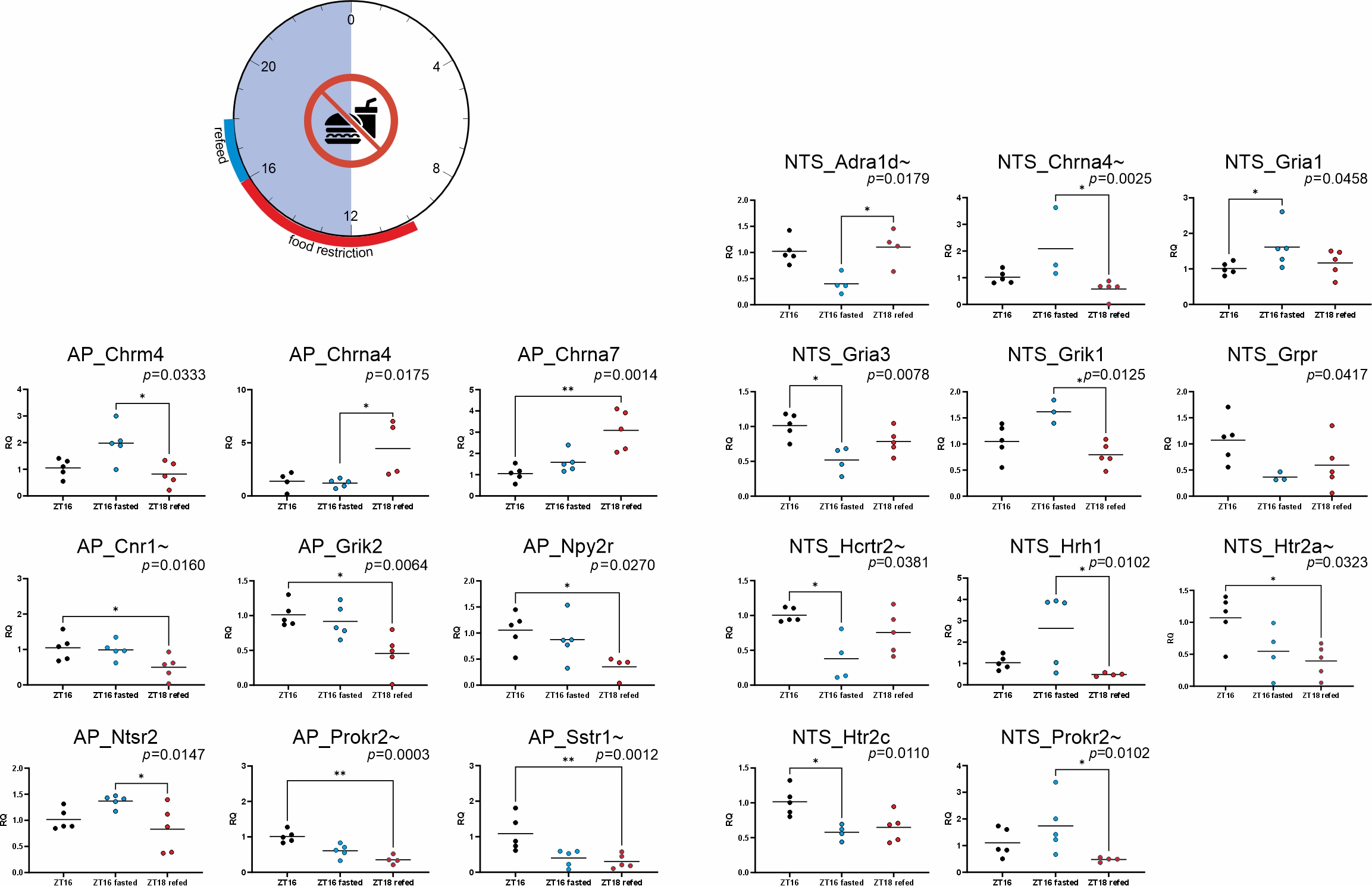

### Supplemental Figure 5

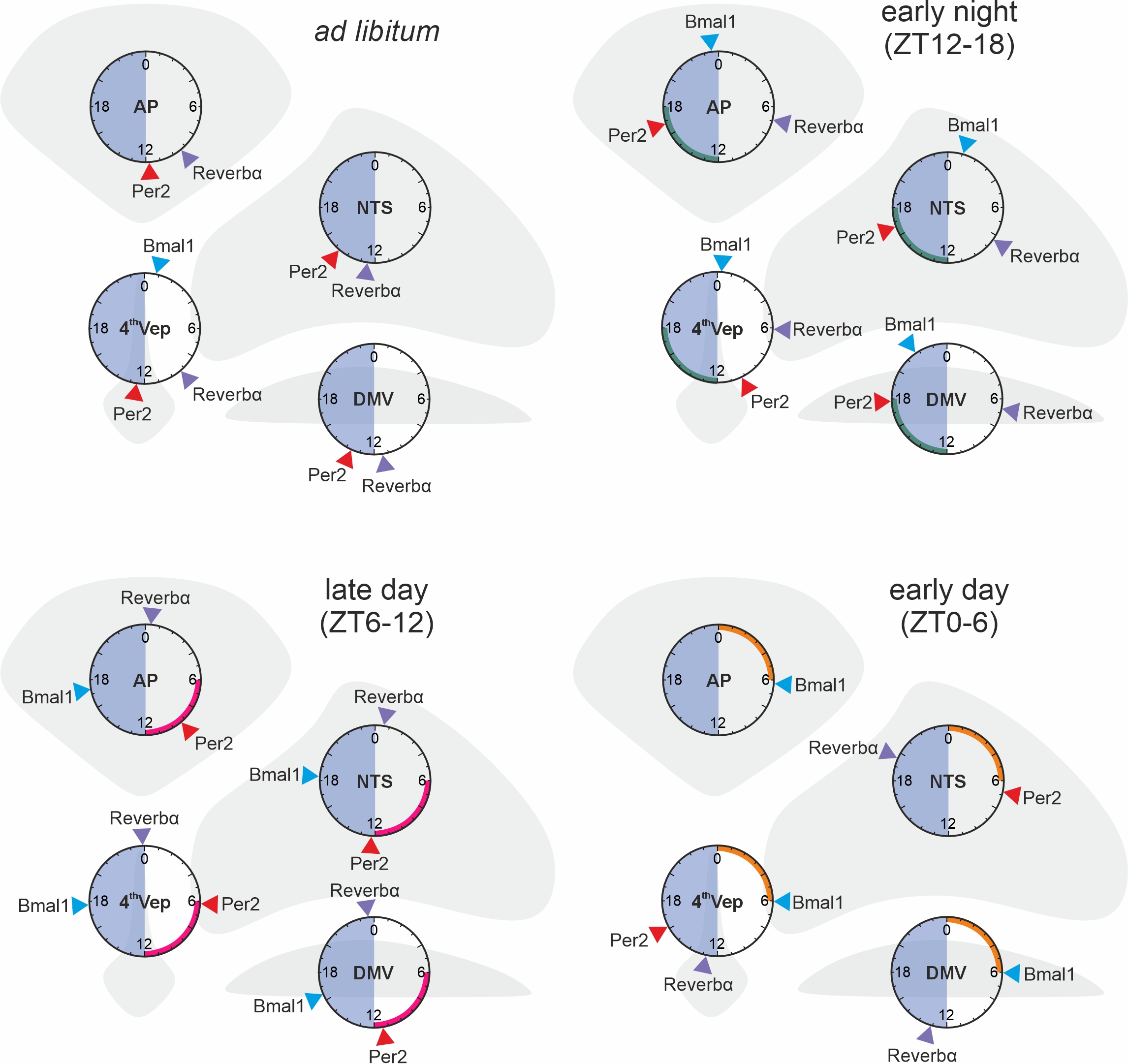

### Supplemental Figure 6

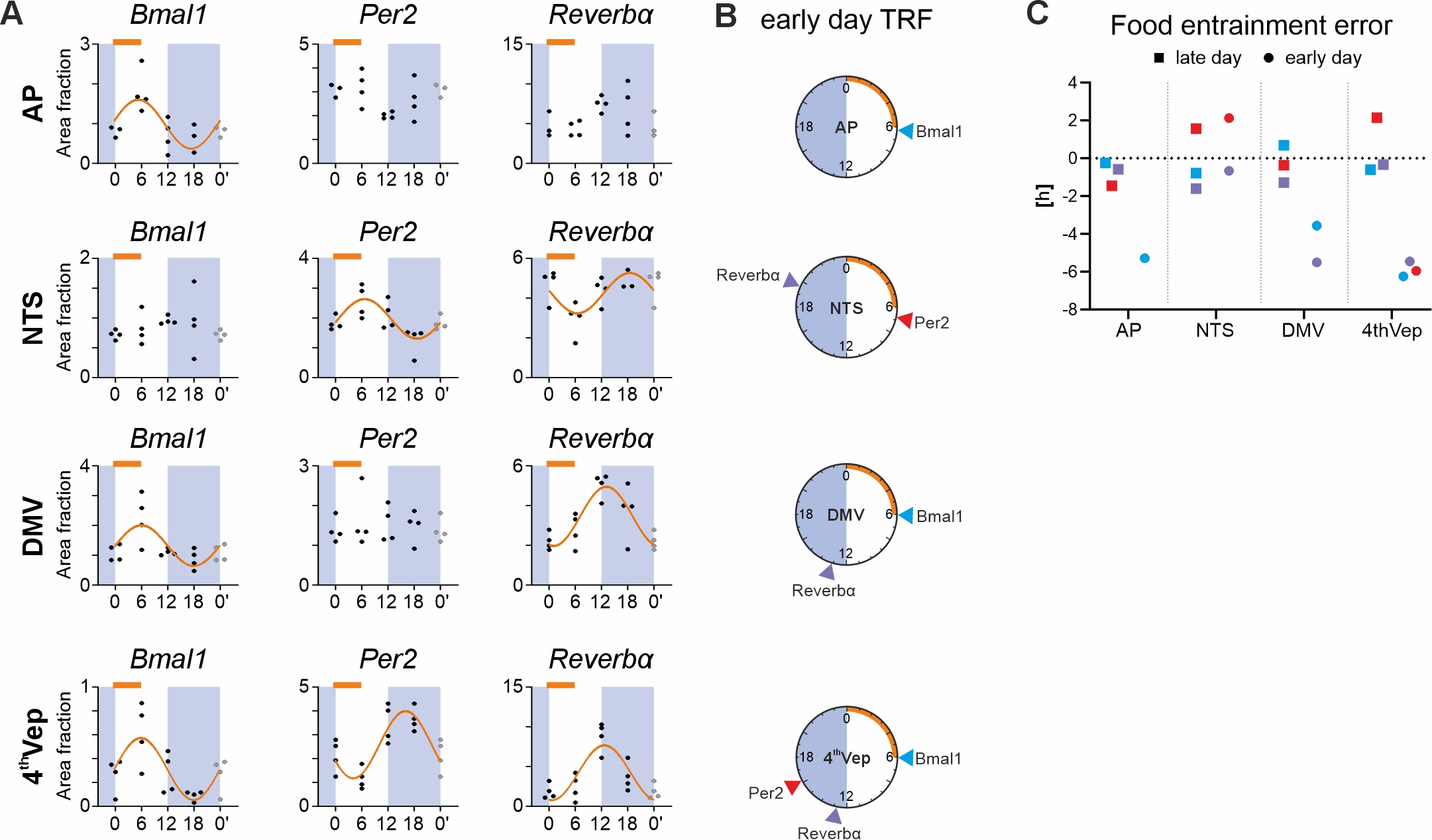
